# Cell shape independent FtsZ dynamics in synthetically remodeled cells

**DOI:** 10.1101/335356

**Authors:** Bill Söderström, Alexander Badrutdinov, Helena Chan, Ulf Skoglund

## Abstract

The FtsZ protein is a key regulator of bacterial cell division. It has been implicated in acting as a scaffolding protein for other division proteins, being a force generator during constriction, and more recently, as an active regulator of septal cell wall production. During an early stage of the division cycle, FtsZ assembles into a heterogeneous structure coined the “Z-ring” due to its resemblance to a ring confined by the midcell geometry. While *in vitro* experiments on supported lipid bilayers have shown that purified FtsZ can self-organize into a swirling ring roughly the diameter of a bacterial cell, it is not known how, and if, membrane curvature affects FtsZ assembly and dynamics *in vivo*.

To establish a framework for examining geometrical influences on proper Z-ring assembly and dynamics, we sculptured *Escherichia coli* cells into unnatural shapes, such as squares and hearts, using division- and cell wall-specific inhibitors in a micro fabrication scheme. This approach allowed us to examine FtsZ behavior in engineered “Z-squares” and “Z-hearts”, and in giant cells up to 50 times their normal volume. Quantification of super-resolution STimulated Emission Depletion (STED) nanoscopy data showed that FtsZ densities in sculptured cells maintained the same dimensions as their wild-type counterparts. Additionally, time-resolved fluorescence measurements revealed that FtsZ dynamics were generally conserved in a wide range of cell shapes. Based on our results, we conclude that the underlying membrane environment is not a deciding factor for FtsZ filament maintenance and treadmilling *in vivo*.

## Introduction

Most bacterial cells divide by binary fission, whereby one mother cell splits into two identical daughters ^1-3^. Decades of study have led to a detailed understanding of how the cell division machinery, the divisome, carries out this task during the later stages of the cell cycle ^4,5^. At the heart of this process is the eukaryotic tubulin homologue, FtsZ ^6^ that, together with its membrane anchors ZipA and FtsA, forms an intermediate structure called the proto-ring (Fig. 1a) ^7^. Functioning as a recruitment base, the proto-ring components then enlist the remaining essential division proteins to form a mature ‘divisome’ ^5^. As soon as it is fully assembled, the divisome starts to constrict the cell envelope by reshaping the septal geometry, ultimately leading to sequential closure of the inner and outer membranes ^8-10^.

**Figure 1.**
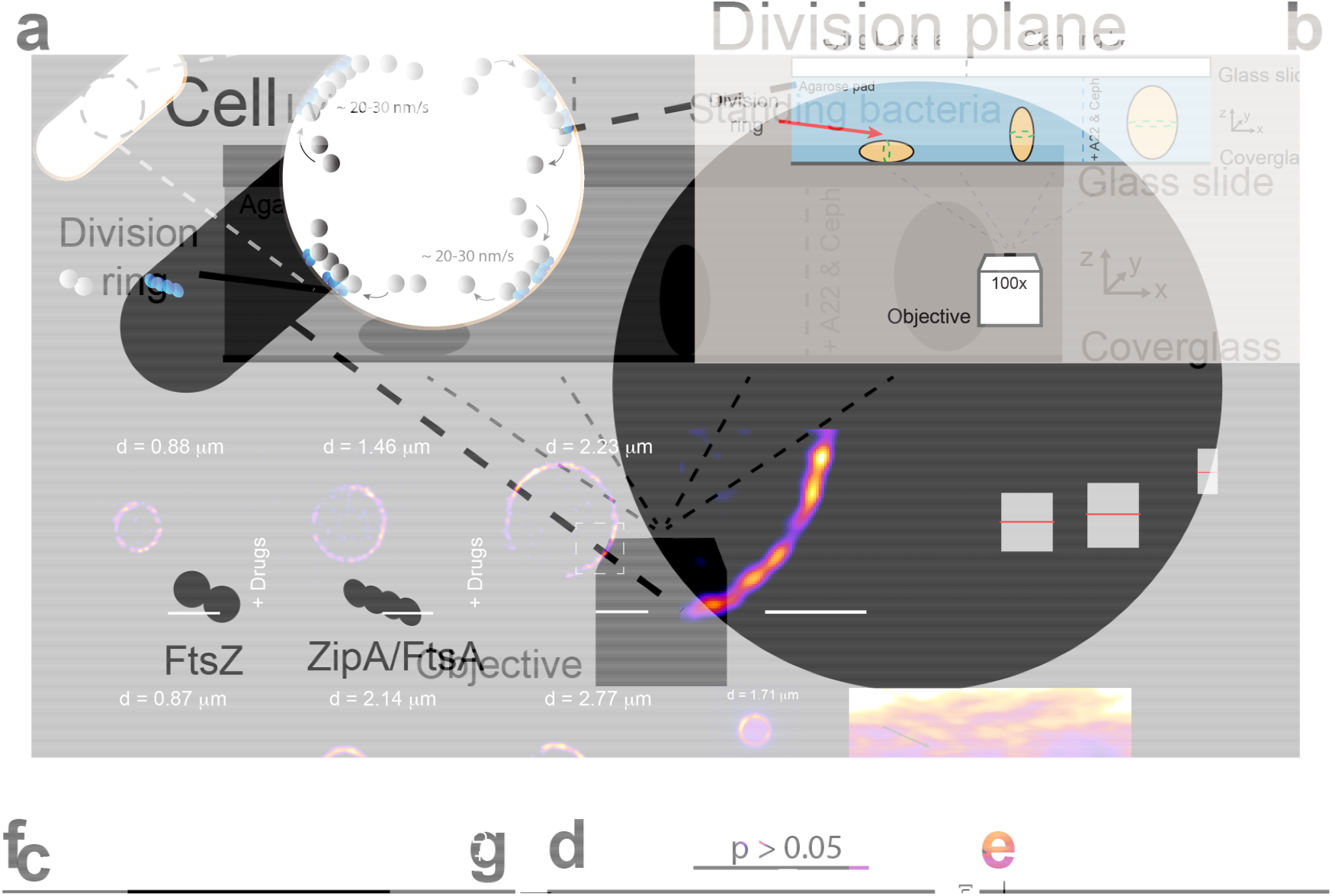
Midcell Z-ring assembly is unaffected by increased cell diameter. **a**, Simplified cartoon showing FtsZ treadmilling at the division plane of an *E. coli* cell. For clarity, only the membrane tethers, FtsA and ZipA, are shown. **b**, Schematic representation of cell placement for imaging. Standing cells were trapped in a vertical position in micron-sized holes in agarose pads created using micron-sized pillars. Conditions for proper division ring placement are met when width < length. The left and middle cells represent untreated cells. The cell on the right has increased dimensions due to drug exposure (A22 and cephalexin). **c**, Time-gated STED (gSTED) image of a typical FtsZ-ring (FtsZ-mNeonGreen) in an untreated standing cell. Scale bar = 1 μm. **d** and **e**, gSTED images of FtsZ-mNeonGreen rings in *E. coli* cells treated with drugs, showing increased ring diameter. Scale bar = 1 μm. “Drugs” refer to A22 and cephalexin. **f**, Close-up of representative FtsZ densities shown in **e**, from a cell with increased diameter. Scale bar = 0.5 μm. **g**, Quantification of FtsZ density lengths in untreated and drug-treated cells. Mean ± S.D. was 122.8 ±?43.9 nm (n = 77) and 132.4 ± 48.7 nm (n = 172) for untreated and drug-treated cells, respectively. No statistically significant difference was measured, p > 0.05. Inset shows density widths in drug-treated cells, mean ± S.D. = 88.4 ± 9.8 nm (n = 172). **H - k**, Structured Illumination Microscopy (SIM) images of FtsZ-GFP in *E. coli* cells (**h**) untreated or (**i** - **k**) treated with drugs. Scale bars = 1 μm. **l**, Snapshots of epifluorescence (EPI) images from time-lapse series of FtsZ-GFP dynamics in drug-treated cells. Scale bars = 1 μm. Corresponding kymographs are shown adjacent to each image. Black arrows point to examples of FtsZ trajectories. **m**, Average treadmilling speed of FtsZ-GFP in untreated (26 ± 15 nm/s, n = 102) and drug-treated cells (30 ± 18 nm/s, n = 102). “d” in (**c** - **e**) and (**h** - **l**) indicates cell diameter.

In rod-shaped model bacteria such as *Escherichia coli* and *Bacillus subtilis*, FtsZ is believed to organize into short bundles of filaments, roughly 100 nm in length ^11,12^, that treadmill at the septum with a circumferential velocity in the order of 20-30 nm/s ^13-15^. The treadmilling filaments guide and regulate septal peptidoglycan (PG-) production and ingrowth, leading up to septation ^16^. This mode of action may be limited to rod-shaped bacteria that have two separate PG-machineries, as opposed to cocci, which have only one PG-machinery that is capable of finalizing division in cells with inhibited FtsZ dynamics ^17^.

At a late stage of membrane constriction, but prior to inner membrane fusion, FtsZ disassembles from midcell, indicating the possible existence of an upper limit of ring curvature ^6,9^. However, other geometrical constraints that might govern Z-ring maintenance and stability are currently unclear. We were curious as to whether geometrical changes to cell shape would influence Z-ring formation and dynamics. In this study, we examined FtsZ formation, organization and behavior in *E. coli* cells that were sculptured into complex geometrical shapes in micron sized holes.

## Results

### FtsZ structure and dynamics in Z-rings are not sensitive to increased ring size

As a reference for unmodified division rings, we imaged Z-rings in *E. coli* cells expressing FtsZ-mNeonGreen as the only source of FtsZ ^18^. Under our experimental conditions, this strain produced normal-looking, sharp Z-rings (Supplementary Fig. S1) and grew and divided similarly to wild-type (WT) *E. coli* (MC4100) (Supplementary Fig. S2a-e). We then trapped the cells in a vertical position in micron-sized holes that were produced in agarose pads using silica micron pillar arrays ^14^ (Fig. 1b, Supplementary Fig. S3), and imaged the cells using super-resolution time-gated Stimulated Emission Depletion (gSTED) nanoscopy. In these standing cells, a heterogeneous Z-ring with distinct FtsZ-mNeonGreen densities was clearly seen traversing the circumference of the cell (Fig. 1c), similar to what has been observed before ^12,14^.

Previous work has shown that FtsZ densities generally maintain the same length throughout envelope constriction ^12,14^. We wanted to see if this was also true for cells growing in the opposite direction, *i.e.* would FtsZ densities maintain the same dimensions in Z-rings of cells with increased diameter at midcell? In order to increase cell diameter, we treated *E. coli* cells with A22 and cephalexin (hereafter collectively referred to as ‘drugs’), in a way similar to what has previously proven successful for cell shape manipulations ^19^. A22 disrupts MreB dynamics and therefore perturbs the characteristic rod-shape of *E. coli* cells ^20,21^, while cephalexin blocks cell division by inhibiting the transpeptidase activity of FtsI ^22^. The net effect of this dual drug treatment is the growth of cells into shapeable blebs that are unable to divide (Supplementary Fig. S4a).

As long as cell width remains less than cell length, FtsZ molecules should be directed to midcell by the Min system ^19^, such that a ring-like structure should be observed in the xy-plane of vertically-oriented, standing cells (Fig. 1b). To confirm this, we exposed *E. coli* cells expressing FtsZ-mNeonGreen to drugs, and then trapped the cells vertically in holes with a depth of 4.5 - 6 μm, and a diameter of up to 3.5 μm. Depending on the size of the holes, cells were incubated between 120 and 240 minutes prior to imaging; over-incubation resulted in cells that outgrew the holes (Supplementary Fig. S4b. Letting cells grow for long time (> 10 h) produced giant blobs with internalized FtsZ-mNeonGreen chain, see SI Text). We found “normal-looking” Z-rings that spanned the midcell circumference for the entire range of cell diameters that were imaged (∼ 1 - 3 μm) (Fig. 1d-e). Importantly, confocal Z-stacks showed that each cell contained only one Z-ring (Supplementary Fig. S5 and Movie SM1). Close inspection of STED images revealed that the Z-rings in larger cells were composed of fluorescent densities (Fig. 1f) with average lengths and widths of 132 ± 48 nm and 88 ± 9 (n = 172), respectively, which were similar (p > 0.05) to Z-ring densities in untreated cells (Fig. 1g).

After we had established that large Z-rings can form in cells with increased diameter, we proceeded to calculate FtsZ dynamics in these larger rings. However, strains expressing FtsZ-FP as the only source of FtsZ have been shown to have a phenotype similar to that of FtsZ mutants deficient in GTPase activity, with severely impaired treadmilling speed ^13^. Therefore, we chose to image cells that expressed FtsZ-GFP from an ectopic locus on the chromosome, in addition to native FtsZ ^23^. Earlier studies showed that FtsZ-GFP, when expressed at levels below 50 % of total cellular FtsZ levels, caused no observable phenotypic changes ^9,12,23,24^. In our experimental setup, FtsZ-GFP was expressed at ∼ 30 % of total FtsZ levels (Supplementary Fig. S2).

Structured Illumination Microscopy (SIM) of drug-treated *E. coli* cells expressing FtsZ-GFP showed large heterogeneous rings that were similar to those of FtsZ-mNeonGreen (Fig. 1h - k). Time-lapse images revealed that FtsZ densities moved around the midcell circumference, even in Z-rings with a diameter up to three times larger than that of a WT cell (Supplementary Movie SM2). There was no difference in the speed of individual densities in the rings of untreated cells compared to those in sculptured cells that had a diameter 50 - 200 % larger than WT (26 ± 15 nm/s and 30 ± 18 nm/s, respectively) (Fig. 1l - m), suggesting that filament treadmilling speed is not influenced by the length of the cell circumference. ZipA-GFP, an FtsZ membrane anchor, also moved at essentially the same speed in both normal- and large-sized rings (26 ± 8 nm/s) (Supplementary Fig. S6 and Movie SM3), which is comparable to previously reported speeds ^14^.

Since treadmilling behavior of FtsZ in large cells was very similar to that in WT cells, we were curious to see whether FtsZ subunit exchange in the rings would also be similar. To assess this, we performed Fluorescence Recovery After Photobleaching (FRAP) experiments on both untreated and drug-treated cells. We bleached half of the FtsZ-GFP molecules in the rings of standing cells and monitored fluorescence recovery over time (Fig. 2a). Z-rings in untreated cells had a mean t_1/2_ recovery time of 8.4 ± 1.9 sec, n = 23 (Fig. 2b), consistent with previous studies ^14,25^. Surprisingly, the average t_1/2_ recovery time was the same for Z-rings with a wide range of diameters (Fig. 2b). We believe this reflects the formation of a greater number of FtsZ filaments at the inner membrane of expanded cells, due to larger accessible surface area (Fig. 2c), while filament treadmilling speed remains unchanged.

**Figure 2.**
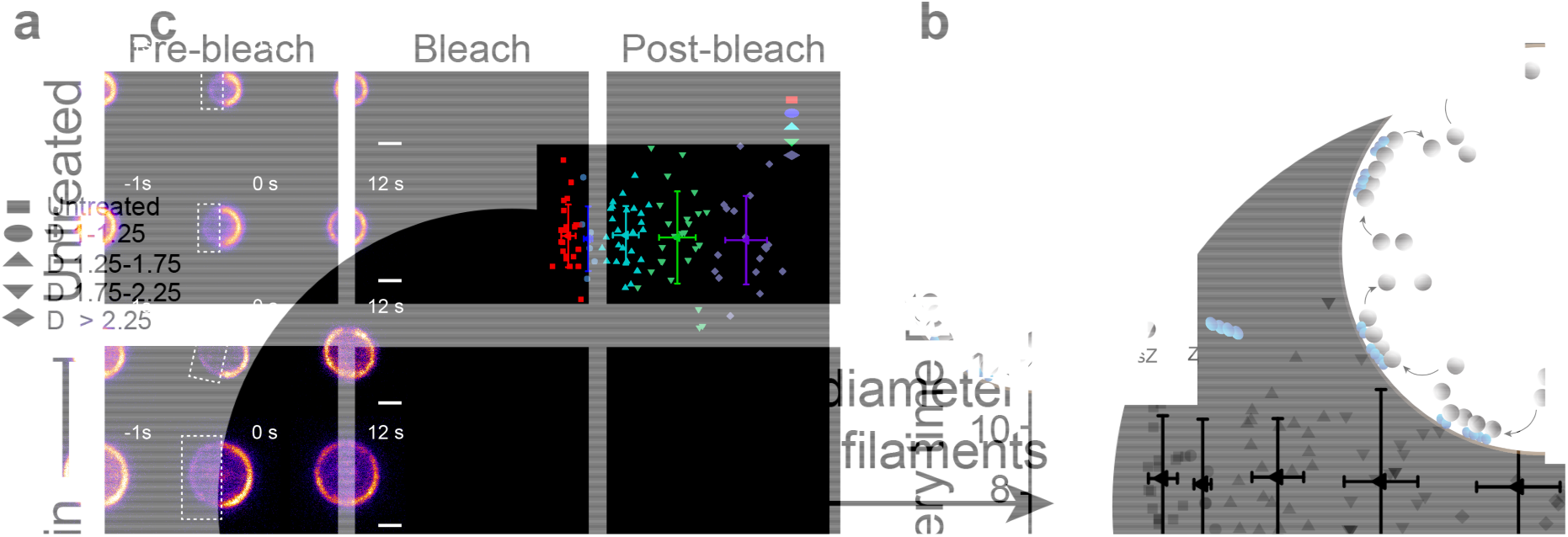
Cell size independent recovery of fluorescence in FtsZ-rings. FRAP measurements on FtsZ-GFP rings in *E. coli* cells trapped standing in a vertical position. **a**, Representative cells of different diameter, untreated or treated with drugs. White boxes indicate bleached areas. Scale bars = 1 μm. **b**, Quantification of FRAP data from untreated cells (red), and drug-treated cells (other colors), showing that fluorescence recovery time is independent of cell diameter (in the range investigated, *i.e.* ∼ 1 - 3 μm). n_tot_ = 105. “D” = cell diameter range. (see SI text for detailed values). **c**, Schematic representation of possible FtsZ filament distribution in cells of different diameters. Cells with larger diameter can accommodate a greater number of FtsZ filaments.

### The ‘FtsZ-square’

Next, we wanted to know if drug-treated cells placed in deep (5 μm) rectangular volumes would adapt to these shapes and effectively form ‘Z-rectangles’ or ‘Z-squares’ instead of ‘Z-rings’. Previous work has shown that cells can adapt to rectangular shapes in shallow wells, approximately 1 μm deep ^19^. Here, we produced quadrilateral patterns in agarose pads using silica micron pillar arrays similar to those previously described ^14^, with the exception that the pillars were rectangular and 5.5 ± 0.5 μm in height. Side lengths of the micron chambers were up to 3.5 μm in length (Supplementary Fig. S7), resulting in well volumes up to 80 μm^3^, roughly 50-fold larger than the volume of a WT cell (assuming a WT cell size of 2 μm in length and 1 μm in width) (Supplementary Fig. S8).

Drug-exposed cells expressing FtsZ-mNeonGreen were placed in rectangular micron holes and incubated at room temperature for 300 - 420 minutes (longer incubation times were needed due to increased well size). The cells adapted to their new shapes and formed rectangular cuboids with only one “Z-square” per cell (Fig. 3a, Supplementary Movie SM4). Notably, FtsZ densities were observed both in the sharp corners and along the sides of the rectangles (Fig. 3b, Supplementary Fig. S9). Quantification of the FtsZ-mNeonGreen densities showed that they had similar dimensions to those in untreated cells, with an average length of 105.4 ± 39.6 nm and width of 79.6 ± 18.2 nm (n = 147) (Fig. 3c). This suggests that FtsZ filament dimensions *in vivo* are insensitive to membrane curvature (or lack thereof).

**Figure 3.**
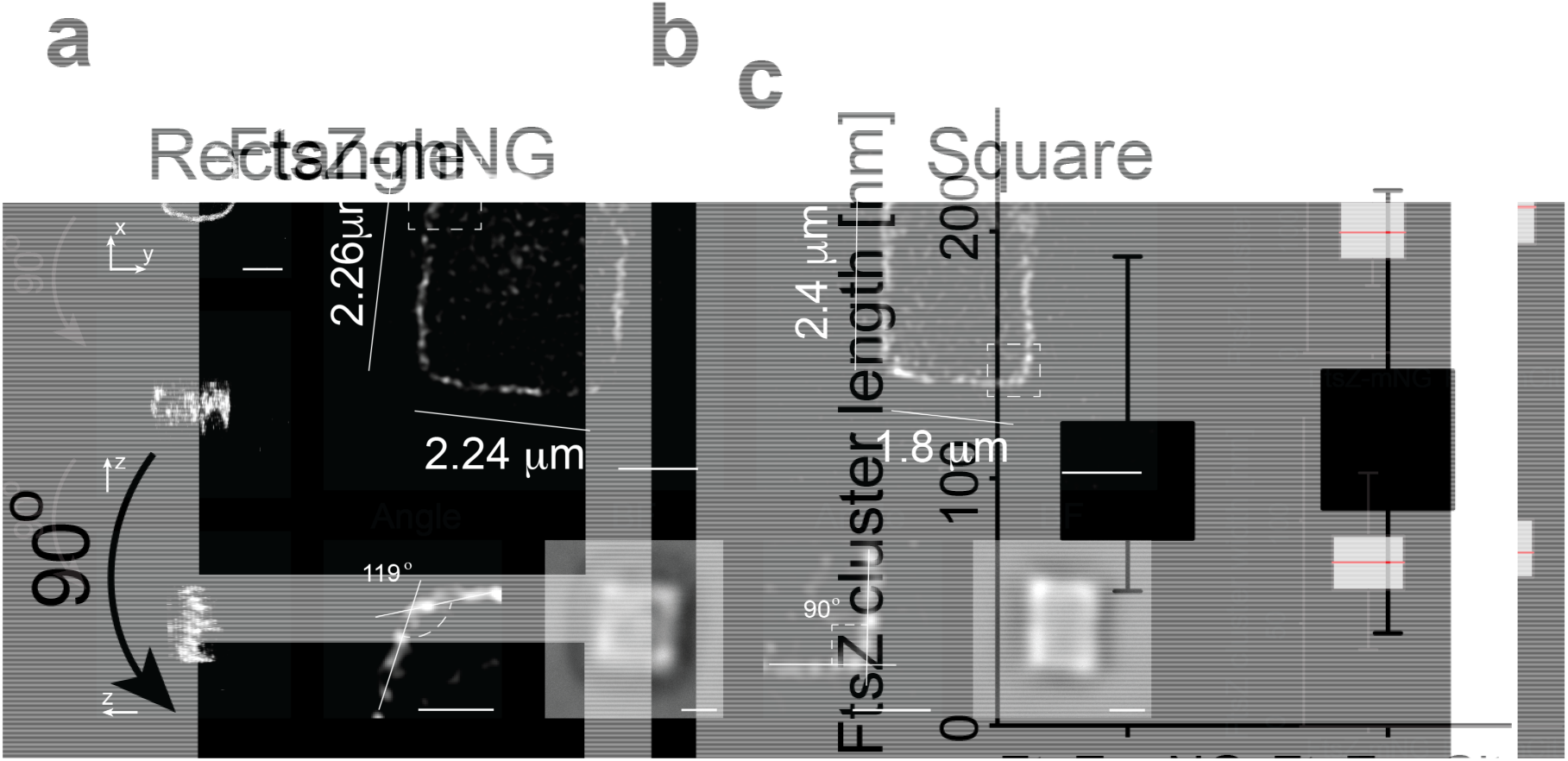
FtsZ-squares and -rectangles in shaped cells. Drug-treated (A22 and cephalexin) *E. coli* cells expressing FtsZ-mNeonGreen were sculptured into rectangular shapes and imaged using super-resolution STED nanoscopy. **a**, 3D rendering of a confocal Z-stack of an FtsZ-mNeonGreen square, showing only one band of FtsZ. Note that information along the z-axis is elongated. **b**, Representative STED images of FtsZ-mNeonGreen in square (left) and rectangular (right) cells with perimeters ranging from 8.4 to 11.52 μm (compared to WT ∼ 3 μm). Additional examples are provided in Supplementary Fig. S9. Close-up images show representative corner angles. BF, brightfield image of corresponding cells. **c**, Quantification of FtsZ cluster dimensions, showing little difference between FtsZ-mNeonGreen (105.4 ± 39.6, 79.6 ± 18.2; length and width, respectively. n = 147) and FtsZ-mCitrine (118.3 ± 41.3, 86.3 ± 22.5; length and width, respectively. n = 162. Example images of FtsZ-mCitrine squares are shown in Supplementary Fig. S10). Scale bars = 1 μm.

To generate a fluorescent FtsZ fusion protein that could be used for both super-resolution STED imaging and examination of filament dynamics when grown in rich media at 37 °C, we constructed a plasmid-expressed FtsZ-mCitrine fusion. FtsZ-mCitrine was expressed from an IPTG-inducible, medium copy-number plasmid, pTrc99a, at a level approximately equal to 30 % of total cellular FtsZ. Under these conditions, FtsZ-mCitrine formed normal-looking, sharp Z-rings (Supplementary Figs. S1 and S2). Cells expressing FtsZ-mCitrine were then exposed to drugs, trapped in rectangular micron-sized holes, and incubated for 180 - 280 minutes at room temperature before gSTED imaging. We found that FtsZ-mCitrine formed filaments that were 118.3 ± 41.3 nm long and 86.3 ± 22.5 nm wide (n = 162), similar to FtsZ-mNeonGreen filament dimensions (Fig. 3c, Supplementary Fig. S10), indicating that fluorophore choice did not influence cluster dimensions in the rings. For consistency, we also imaged rectangular cells expressing FtsZ-GFP from the chromosome using SIM (Supplementary Fig. S10). All three strains tested adapted to the rectangular shape, producing sharped-cornered “Z-rectangles”.

### FtsZ dynamics in rectangular-shaped cells

In order to examine the dynamics of FtsZ in rectangular cells, we performed time-lapse imaging on cells expressing either FtsZ-mCitrine or FtsZ-GFP. Although a few fluorescence spots were abnormally bright and immobile (∼ 1 spot / 5 cells, with a maximum of 2 spots in one cell) (Fig. 4b, Supplementary Movie SM7. Red arrow), the majority of FtsZ densities were highly dynamic (Fig. 4a-b, Supplementary Movies SM5 - SM6). Note that the bright, immobile spots were excluded from treadmilling analyses. Close inspection of time-lapse sequences suggested that FtsZ bundles in rectangular-shaped cells may be able to treadmill, even in right-angled corners (Fig. 4b-c, Supplementary Movie SM8). The average speed of FtsZ-mCitrine densities in rectangular cells with perimeter lengths up to 13 μm (more than four times the circumference of a WT cell) was 27.6 ± 12.5 nm/s (n = 109), which was consistent with the measured treadmilling speed of FtsZ-GFP in rectangular cells (25.3 ± 11.3 nm/s, n = 122) (Fig. 4d), large cylindrical cells (30 ± 18 nm/s, Fig. 1m) and untreated cells (∼ 25 nm/s) ^13,14^.

**Figure 4.**
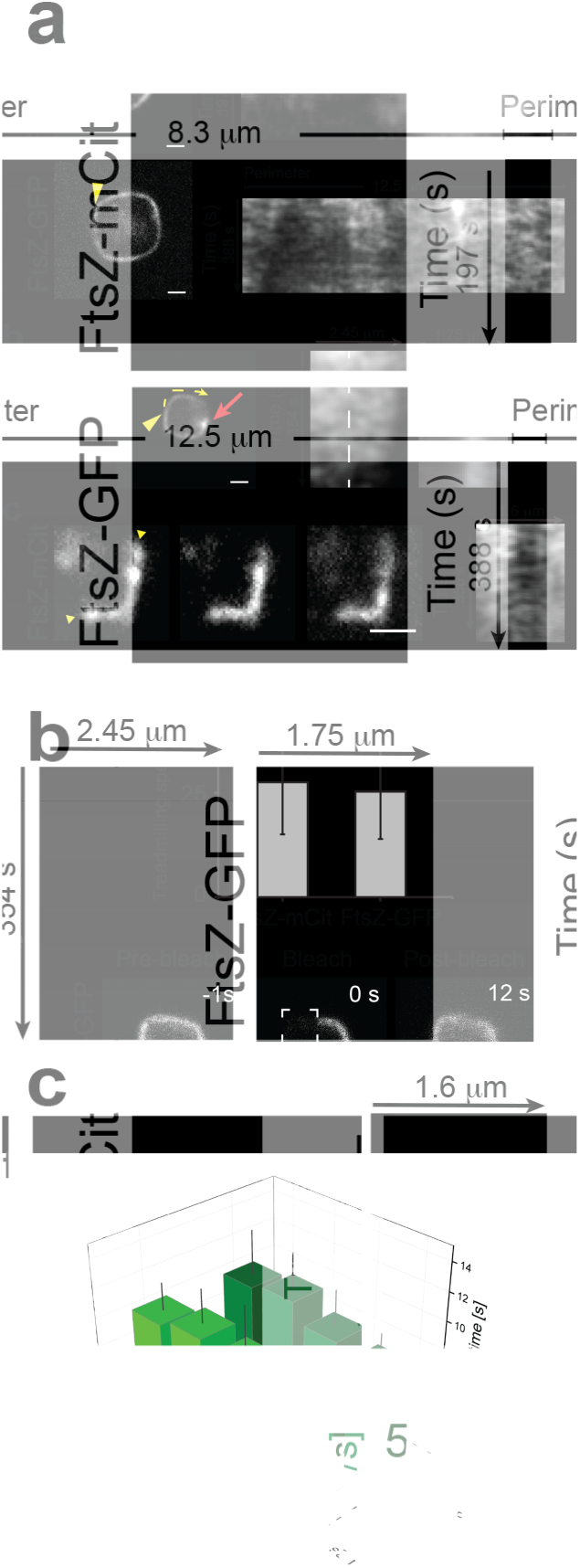
FtsZ dynamics in rectangular-shaped cells. The dynamics of FtsZ in rectangular shapes were assessed by time-lapse imaging and FRAP measurements on *E. coli* cells expressing FtsZ-mCitrine or FtsZ-GFP. **a** - **c**, Snapshot images from time-lapse series of FtsZ-mCitrine or FtsZ-GFP in rectangular shaped cells. Corresponding kymographs are shown next to each image. **a**, Kymographs were taken around the entire perimeter (starting in the upper left corner, moving counter-clockwise, indicated by the yellow arrowheads). **b**, Kymographs were taken along the yellow line starting at the yellow arrowhead (left kymograph), or over the bright spot indicated by the red arrow (right kymograph). The white striped line in **b** indicates the upper left corner of the cell. **c**, Kymograph taken between the yellow arrowheads (top to bottom is left to right in the kymograph). **d**, Average treadmilling speed of FtsZ-mCitrine and FtsZ-GFP in rectangles was 27.6 ±?12.5 nm/s (n = 97) and 25.3 ± 11.3 nm/s (n = 122), respectively. **e**, Typical FRAP measurement of FtsZ-GFP in a rectangular *E. coli* cell. Half of the rectangle was bleached. **f**, Average recovery times for FtsZ-mCitrine (light, n_tot_ = 24) and FtsZ-GFP (dark, n_tot_ = 22) in FtsZ-rectangles of various perimeter lengths (see SI text for detailed values). Scale bars = 1 μm.

To determine whether the dynamics of FtsZ subunit exchange are affected by changes to circumferential length and shape, we collected FRAP measurements on FtsZ bundles in rectangular-shaped cells (Fig. 4e, Supplementary Movie SM9). The recovery times of half-bleached rectangles of varying sizes matched those of rings, with mean t_1/2_ recovery times of 9.85 ± 2.58 s (n = 24) and 9.15 ± 2.55 s (n = 22) for FtsZ-mCitrine and FtsZ-GFP, respectively (Fig. 4f). This suggests that subunit exchange from the cytoplasmic FtsZ pool is independent of circumference length and membrane curvature. The data thus far indicate that the maintenance and dynamics of FtsZ filaments are preserved in both large Z-rings and Z-rectangles of varying size.

### FtsZ dimensions and dynamics in heart-shaped cells

To examine whether FtsZ could literally be (at) the heart of cell division, we engineered micron pillar arrays that were heart-shaped (Supplementary Fig. S11). Heart shapes were chosen because they would sculpt cells in such a way that highly curved, straight, and angled membrane segments would be present within a single cell. Drug-treated *E. coli* cells expressing cytoplasmic GFP, FtsZ-mNeonGreen or FtsZ-mCitrine were sculptured into hearts as described above (Fig. 5a). Perhaps not surprisingly, quantification of 155 individual FtsZ densities from the heart-shaped cells revealed dimensions similar to those in round and rectangular cells (129 ± 44 nm long and 84 ± 9 nm wide) (Fig. 5b). We also found that the average speed of FtsZ-mCitrine in heart-shaped cells (22 ± 10 nm/s, n = 44) was essentially the same as that in untreated cells (Figure 5c, Supplementary Movie SM10).

**Figure 5.**
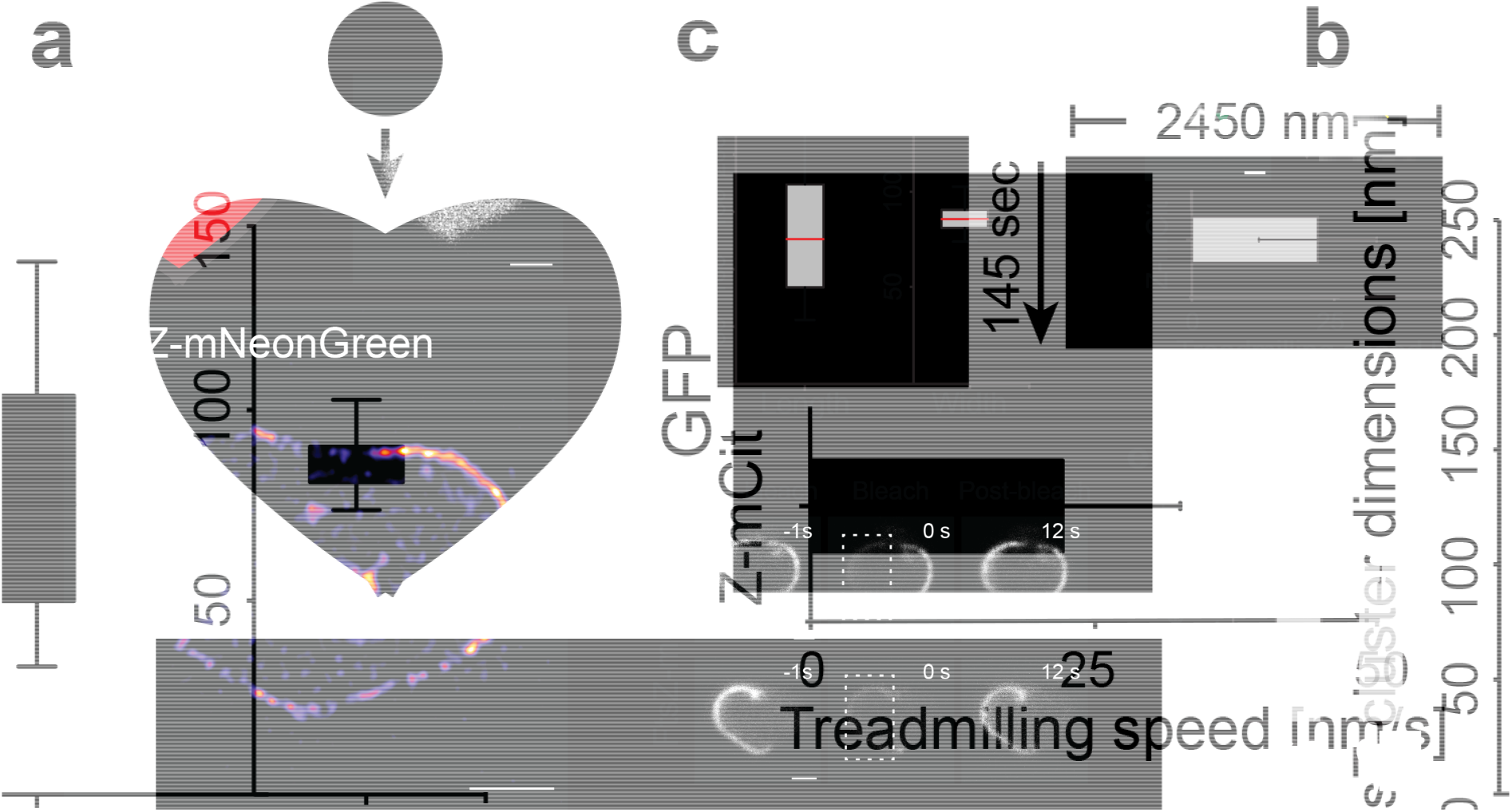
FtsZ cluster dimensions and dynamics in heart-shaped cells. FtsZ behavior in *E. coli* cells sculptured into heart shapes. **a**, Upper left, Cartoon representation of a WT *E. coli* cell and a heart shape, highlighting the large and complex structural changes of a cell-to-heart transition, approximately to scale. Upper right, Drug-treated cell expressing cytoplasmic GFP, shaped as a heart. Lower, STED image of an “FtsZ-heart” (FtsZ-mNeonGreen) in a drug-treated *E. coli* cell. **b**, Lengths and widths of 155 individual FtsZ-mNeonGreen fluorescence densities in cells shaped as hearts. Average length = 129 ± 44 nm and width = 84 ± 9 nm. **c**, Upper row, SIM image from a time-lapse series (epi-fluorescence) of a heart-shaped cell expressing FtsZ-mCitrine. Green arrowhead indicates internal FtsZ clustering. Corresponding kymograph is shown adjacent to the image, and was generated starting at the yellow arrowhead in the SIM image, moving counter-clockwise for the indicated length. The yellow arrow points to an FtsZ trajectory. Lower, average treadmilling speed of FtsZ-mCitrine (Z-mCit) filaments in hearts (22.6 ± 10.4 nm/s, n = 44). **d**, FRAP measurements of FtsZ-mCitrine in heart-shaped cells. Top row, bleaching of half the FtsZ-mCitrine molecules in a ‘full’ heart. Bottom row, bleaching of a ‘half-full’ heart. No difference in recovery time was observed. **e**, Histogram of average t_1/2_ recovery times calculated from FRAP measurements. Recovery in ‘full hearts’: 7.1 ± 1.1 s (n = 24), recovery in ‘half hearts’: 6.9 ± 0.9 s (n = 9).

For about one-third of the heart-shaped cells, we noticed bright spots of internalized FtsZ-FP signal that accumulated close to the cell center (Figure 5c, green arrowhead). Although we couldn’t distinguish whether these were true FtsZ clusters or aggregated protein, cytoplasmic clustering of FtsZ in WT cells have previously been reported ^12^. Furthermore, although most hearts had FtsZ-FP signal spanning the full perimeter of the cell, approximately 20 % were only “half full” (Fig. 5d, left). We do not fully understand the underlying reason for this, however it is unlikely due to image focus or cell tilt issues, as every cell was scanned in the z direction prior to imaging. Nevertheless, when we subjected the heart-shaped cells to FRAP, fluorescence recovery rates were equal for both full and half-full hearts (Fig. 5d), with mean t_1/2_ recovery times of 7.1 ± 1.1 s (n = 24) and 6.9 ± 0.9 s (n = 9), respectively (Fig. 5e).

### FtsZ-“rings” form in complex cell shapes

To explore if cell geometry plays a role in Z-“ring” formation, we set out to remodel cells into other complex shapes. Even though highly complex-shaped bacteria occur in nature, such as star-shaped bacteria ^26^, we wanted to test whether rod-shaped *E. coli* cells would allow themselves to be drastically remodeled. Using micron pillars of various shapes, we produced holes in agarose pads such that drug-exposed cells could be sculptured into complex shapes, such as pentagons, half-moons, stars, triangles and crosses (Fig. 6a, middle row. Supplementary Fig. S11). The cells conformed remarkably well to these shapes, forming sharp boundary angles < 70° (Fig. 6a, Star). After we confirmed that cells could adapt to these complex shapes, we placed cells expressing FtsZ-mCitrine into the micron holes, allowed for reshaping to occur, and then imaged the cells using STED nanoscopy. Cells of all tested shapes produced easily recognizable FtsZ-“shapes” at midcell (Fig. 6a, bottom row). Subsequent analysis of the lengths and widths of the FtsZ densities revealed little difference in dimensions between the different shapes, suggesting a minimal role of cell shape in determining FtsZ cluster dimensions *in vivo* (Fig. 6b). Additionally, time-lapse imaging of cells expressing FtsZ-mCitrine in various shapes showed similar dynamics to those measured in untreated cells (Supplementary Movie SM10).

**Figure 6.**
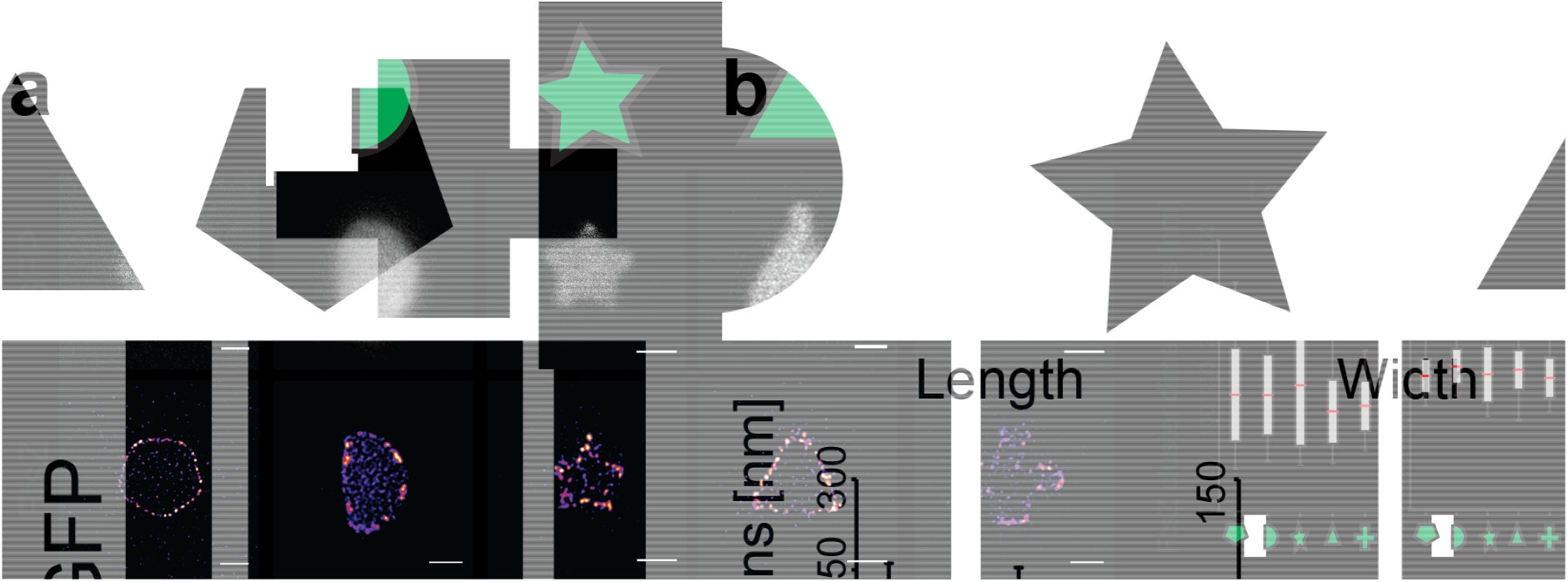
FtsZ bundle dimensions in complex shapes. **a**, Cells expressing cytosolic GFP or FtsZ-mCitrine were remodeled into various shapes. Top row, schematic representation of the cell shapes. Middle row, representative cells expressing cytosolic GFP, and sculptured in the corresponding shapes. Bottom row, an FtsZ-pentagon, FtsZ-half-moon, FtsZ-star, FtsZ-triangle and FtsZ-cross in sculptured cells. Scale bars = 1 μm. **b**, Quantification of FtsZ densities by length and width in shaped cells (see SI text for detailed values).

## Conclusions

Cells, both bacterial and eukaryotic, have the ability to adapt remarkably well to their local environments ^27−31^, reverting to their original shapes after stress ^32,33^ and dividing with striking midcell accuracy even when remodeled into irregular cell shapes ^27,30^. In bacteria, the tubulin homologue FtsZ assembles into a ring-like structure at midcell and is responsible for overall maintenance of the cell division machinery ^5,6^. The general dynamics and organization of the FtsZ-ring have been shown to be quite similar across many bacterial species ^11,13−15,17,34−37^. Common to these species is confinement of the FtsZ-ring to a circular geometry at midcell. Strikingly, when purified FtsZ (together with its membrane anchor FtsA) is placed on supported lipid bilayers, it assembles into a dynamic, swirling ring-like assembly with a diameter resembling that of wild-type *E. coli* cells (approximately 1 μm), hinting at an intrinsically preferred FtsZ-ring curvature ^6,38^.

In this study, we characterized FtsZ midcell accumulation and dynamics in cell shape-determining environments by ‘looking through the Z-ring’ along the long-axis of cells. We observed normal-looking FtsZ-rings in cells with diameters three times the size found in WT cells. However, this might not be surprising, considering only ∼ 30 % of the pool of FtsZ molecules are in the ring of WT cells at any given point in time ^39^. Quantification of FtsZ dimensions revealed little variation between different cell shapes, such as squares, pentagons, triangles and stars (on average 123 ×; 80 nm, length x width, respectively, and summarized in Table 1), suggesting that local membrane geometry has minimal influence on FtsZ cluster dimensions. Compared to untreated cells, rectangular and heart-shaped cells with perimeter lengths more than four times that of a WT cell exhibited similar overall dynamics of FtsZ, as FtsZ-FP fluorescence densities treadmilled at the same average velocity and FtsZ subunit exchange occurred at similar rates (Table 2), independent of cell shape and size.

**Table 1.**
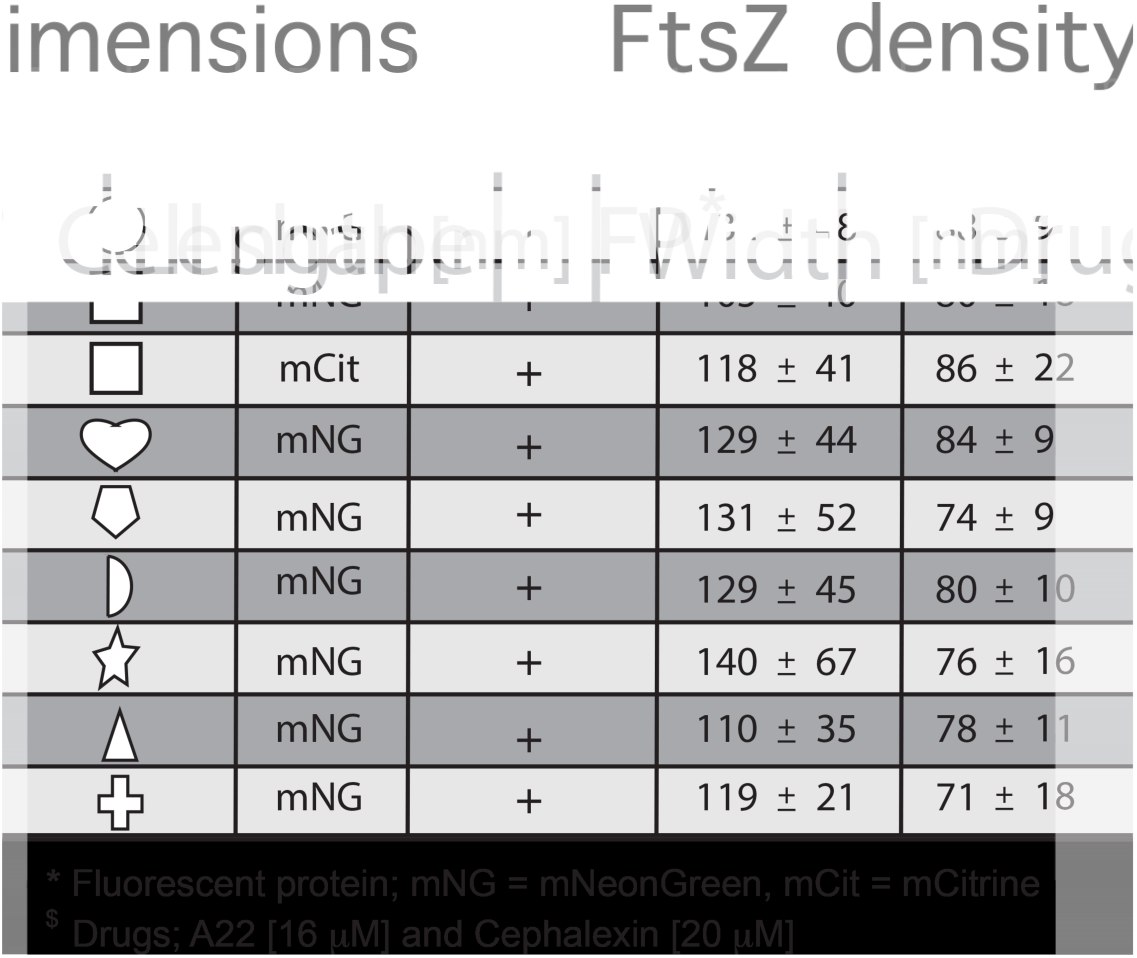
Summary of FtsZ density dimensions at midcell in various cell shapes. In all cell shapes, the average measured density lengths were within 17 % of WT, while average widths were within 13 %. Numbers represent mean ± S.D. Note that values are rounded to whole integers.

**Table 2.**
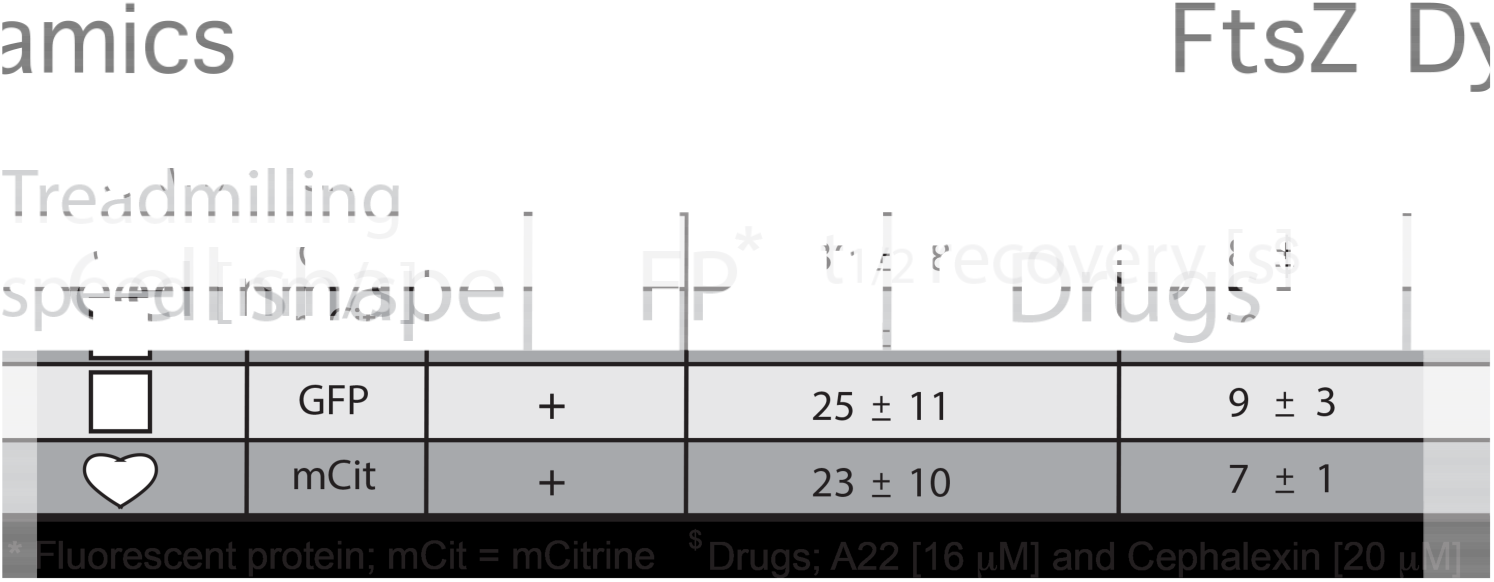
Summary of FtsZ dynamics in various cell shapes. Numbers represent mean ± S.D. Note that values are rounded to whole integers.

In summary, our results from different shaped cells show that Z-“ring” formation and dynamics are not limited to cells of a certain shape or size. This agrees with previous findings, which show that internal cellular structures are maintained in cells that have been reshaped into unnatural forms ^19^. Our observation that FtsZ clusters conform to the geometric shape of the membrane at midcell suggests that FtsZ-ring formation is not affected by changes in membrane curvature. Indeed, cell shape and size are important for proper cellular functions ^40^, however, with the many naturally-occurring shape variations of bacteria ^26,41^, it is perhaps not surprising that FtsZ can adapt to changing environments without compromising its own ability to maintain fundamental functionality. Although our data do not explicitly show that sculptured cells can divide (since downstream division proteins were inhibited), the fact that the dynamic properties of FtsZ are conserved suggests that this may be possible. One particular implication of this is the notion that the Z-ring can be decoupled from the division process but with maintained dynamics, making treadmilling a possible requirement for divisome assembly and organization in rod shaped model bacteria, as previously suggested for cocci ^17^. Presently, we have shown *in vivo* that *E. coli* FtsZ-ring formation and dynamics are conserved, irrespective of cell shape and size.

## Methods

### Bacterial growth

Pre-cultures were grown overnight in 20 ml of rich media (LB) at 37 °C or M9 minimal media supplemented with 1μg ml^−1^ thiamine, 0.2 % (w/v) glucose and 0.1 % (w/v) casamino acids. The following morning, cultures were back-diluted 1:50 in either LB or M9 (with supplements) and antibiotics (ampicillin 25 μg ml^−1^) when needed, and incubated at 30 °C or 37 °C.

### Fluorescent protein production

Chromosomally-encoded FtsZ-mNeonGreen was integrated at the native *ftsZ* locus and did not require any inducer ^18^. Chromosomally-encoded FtsZ-GFP (strain BS001), GFP^CYTO^ (strain BS008) and ZipA-GFP were induced with 2.5 μM, 5 μM and 50 μM IPTG, respectively ^9^.

The plasmid pHC054 (*ftsZ-mCitrine*) was constructed using Gibson assembly ^42^ to generate an IPTG-inducible FtsZ-mCitrine fusion expressed from pTrc99a ^43^. PCR was performed using Q5 High-Fidelity DNA polymerase (New England Biolabs). A DNA fragment containing *ftsZ* was amplified from *E. coli* MC4100 genomic DNA using primers FtsZ(F) (5’-caatttcacacaggaaacagaccatggatgtttgaaccaatggaac-3’) and FtsZ(R) (5’-gcccttgctcaccatctgcaggttgttgttatcagcttgcttacgcagg-3’). *mCitrine* was amplified from mCitrine-N1 plasmid DNA using primers mCitrine(F) (5’-cgtaagcaagctgataacaacaacctgcagatggtgagcaagggcgaggag-3’) and mCitrine(R) (5’-ccgccaaaacagccaagcttttacttgtacagctcgtccatgc-3). pTrc99a plasmid DNA was amplified using primers pTrc99a(F) (5’-ccatggtctgtttcctgtgtg-3’) and pTrc99a(R) (5’-aagcttggctgttttggcgg-3’). The *ftsZ* and *mCitrine* coding regions are separated by a short linker encoding NNNLQ. The plasmid sequence was verified by DNA sequencing (Fasmac, Japan). FtsZ-mCitrine expression was induced with 2.5 μM IPTG. All FtsZ levels were quantified using Western blotting.

### Western blot analysis

Cell extracts from a volume corresponding to 0.1 OD_600_ units were collected for each strain to be analyzed. The extracts were suspended in loading buffer and resolved by SDS-PAGE gel electrophoresis. Proteins were transferred to nitrocellulose membranes using a semi-dry Transfer-Blot apparatus (Bio-Rad). The membranes were blocked in 5 %(w/v) milk and probed with antisera to FtsZ (Agrisera, Sweden) and detected using standard methods.

### Nanofabrication of micro arrays

Micron pillars were engineered using two different, but related, approaches. The first approach was used for round and square/rectangular micron pillars, and was adapted from ^14,15^. Briefly, using a multi-step process similar to that described in ^44^, micron-scale pillars were fabricated on a silicon (Si) substrate by reactive ion etching. A pattern of hard-baked photoresist was created on a Si surface using UV lithography, to work as a mask for etching. Subsequent etching was performed using an Oxford Plasmalab100 ICP180 CVD/Etch system, with a mixture of SF_6_ and O_2_ plasma as an etchant. For our process, a SF_6_:O_2_ ratio of 1:1 was optimal. After etching, the remaining photoresist was removed by O_2_ plasma treatment. Pillar arrays (1 ×; 1 cm or 2 ×; 2 cm) with round pillars were engineered to contain one micron-sized pillar every 5 μm, with dimensions between 0.9 and 3.5 μm wide and 5.25 ± 0.75 μm high (Supplementary Fig. S3). Pillar arrays (1 ×; 1 cm) with square pillars contained micron-sized pillars approximately every 5 μm, with side lengths varying between 1.8 and 3.5 μm, and heights of 5.5 ± 0.5 μm (Supplementary Fig. S7).

To create more complex shapes, a second approach, based on electron beam lithography was used. For this, the micron-scale structures were fabricated on a Si substrate by a multi-step process, which was a combination of electron beam lithography and reactive ion etching techniques. Similar approaches to silicon patterning are described in a number of earlier works ^44−47^. First, a pattern of e-beam resist was created on a Si surface using e-beam lithography. A 50 nm-thick Ti layer was then deposited, and a lift-off process was used to create a metal mask for etching. The use of a metal mask, instead of a baked e-beam resist mask, was necessary due to the high selectivity ratio required for generating structures only a few microns in height. Finally, the etching process was performed as described above, using an Oxford Plasmalab100 ICP180 CVD/Etch system and a mixture of SF_6_ and O_2_ plasma as an etchant. For our process, a SF6:O_2_ flow ratio of 3:2 produced the best results, with a Si:Ti etching selectivity ratio of approximately 100:1. Increased concentration of O_2_ in the mixture has two effects: (i) it improves etching anisotropy, which is essential for avoiding shape distortion from the undercut effect, and (ii) it reduces the selectivity ratio, as the Si etch rate gets slower. After etching, the structures were characterized using a Dektak surface profiler and SEM imaging. The micron structure arrays, which contained various shapes (hearts, triangles, pentagons, half-moons and crosses), were fabricated on 1 ×; 1 cm Si chips with inter-structure distances of approximately 5 μm, and structure heights of 5.5 ± 0.5 μm (Supplementary Fig. S11).

### Micron-sized chamber production and cell growth

Liquefied agarose (5% w/v) in M9 minimal media (supplemented with 0.2 % glucose, 0.1 % casamino acids, 2 μg ml^−1^ thiamine, 40 μM A22 and 20 μg ml^−1^ cephalexin) was dispersed on glass slides and the silica mold (pillar facing downwards) was placed on top. The molds contained either round or rectangular pillars, or various geometrical shapes, as described above. Once the agarose solidified, the mold was removed and ∼ 5 μl of live cell culture at OD_600_ 0.4 - 0.55 (pre-treated with 16 μM A22 for 10 - 15 minutes) was applied on top. To allow the cells to adapt to the different shapes, slides were incubated at RT or 30 °C in a parafilm-sealed petri dish together with a wet tissue to prevent drying. After incubation, cells were covered with a pre-cleaned cover glass (#1.5) for live cell imaging. For STED imaging, cells were first fixed with ice-cold methanol for 5 minutes and carefully rinsed with PBS prior to cover glass application.

### Microscopy

Gated STED (gSTED) images were acquired on a Leica TCS SP8 STED 3X system, using a HC PL Apo 100x oil immersion objective with NA 1.40. Fluorophores were excited using a white excitation laser operated at 488 nm for mNeonGreen and 509 nm for mCitrine. A STED depletion laser line was operated at 592 nm, using a detection time-delay of 0.8 – 1.6 ns for both fluorophores. The total depletion laser intensity was in the order of 20 - 40 MW/cm^2^ for all STED imaging. The final pixel size was 13 nm and scanning speed was either 400 or 600 Hz. The pinhole size was set to 0.9 AU.

Epi-fluorescence and confocal images were acquired on either a Zeiss LSM780 or Zeiss ELYRA PS1 (both equipped with a 100X 1.46NA plan Apo oil immersion objective) with acquisition times between 0.3 and 2 sec. Time-lapse series for generating kymographs were recorded at 2 sec intervals for a time period of at least 118 sec.

SIM images were acquired using a Zeiss ELYRA PS1 equipped with a pco.edge sCMOS camera. The final pixel size in SIM images was 24 nm. Individual images were acquired using an acquisition time of 200 ms per image (a total of 15 images were acquired per SIM image reconstruction) and subsequently reconstructed from the raw data using ZEN2012 software. SIM time-lapse movies (containing at least 14 frames) were recorded without time delays between image stacks.

Confocal Z-stacks (focal plane ± ∼ 3.5 μm) were acquired on a Leica TCS SP8 STED 3X system (operated in confocal mode) using predetermined optimal system settings (Leica, LAS X), with 0.22 μm steps (resulting in 30-32 images per stack), and pinhole size 1 AU. All imaging was performed at RT (∼ 23-24 °C).

### FRAP measurements

Confocal FRAP measurements were performed on a Zeiss LSM780 system using a 100x 1.4 NA plan Apo oil immersion objective and pinhole size 60 μm, as described ^14^. Bleaching was performed for 0.5-0.7 s using 100% laser power applied over the region of interest. Data were collected in time intervals of 1 - 2 sec until steady state was reached. Following background correction, and to account for overall successive bleaching, the fluorescence intensity (F) of the bleached region (half a ring) was normalized to the average ring fluorescence of an unbleached area of the same size, for each time point (t); F_NORM_(t) = F_BLEACHED_(t)/(F_BLEACHED_(t)+ _UNBLEACHED_(t)). All data were exported to Origin9 Pro and data points were fitted to the single exponential function F(t) = F_end_ – (F_end_ – F_start_)* e^−kt^, where F(t) is the fluorescence intensity at time t, F_end_ is the fluorescence intensity at maximum recovery, F_start_ is the fluorescence recovery momentarily after bleaching (at t = 0), and k is a free parameter. The recovery half-time was then extracted from t_1/2_ = ln 2 / k. Importantly, all cells were scanned from top to bottom in order to find the division plane (in which the rings reside).

### Image analysis

Image analysis was performed using Fiji. When necessary, images were background-corrected using a rolling ball with radius 36. Image stacks were motion-corrected using the plug-in StackReg. Kymographs were generated from time-lapse images using the KymoResliceWide plugin (line width 5), from which treadmilling speeds were calculated using the slope of the fluorescence trace, as previously described ^38^.

STED images were deconvolved using Huygens Professional deconvolution software (SVI, the Netherlands). FtsZ-ring diameters were extracted from the average values of the Gaussian fitted fluorescence profiles drawn from 12 - 6 o’clock and 3 - 9 o’clock. Side lengths of shaped cells were determined by applying line profiles in ImageJ. The lengths and widths of individual FtsZ densities were obtained using line scans (line size 4) over at least 5 randomly selected individual fluorescence spots from each deconvolved cell image, whereby a Gaussian was fitted to the intensity profiles in order to extract the Full Width at Half Maximum (FWHM). Note that the long and short axes of each individual FtsZ density were assigned as “length” and “width”, respectively, regardless of orientation relative to the membrane. FtsZ cluster dimensions are given in mean ± S.D. n indicates number of cells, unless explicitly specified.

### Statistical analysis

Boxes represent S.D., with red lines indicating mean. Whiskers on the box plots encompass 95.5% of the distribution. For statistical analyses, two-tailed Student’s *t*-tests were performed using Origin Pro 9. A *p*-value of < 0.05 was considered as statistically significant.

### Data availability

Data presented and material used in this paper can be available upon request from the authors.

## Acknowledgements

The authors would like to thank Harold Erickson (Duke Uni.) for sharing the FtsZ-mNeonGreen strain. Daniel Daley (Stockholm Uni.) is acknowledged for valuable suggestions to improve the manuscript. BS is supported by JSPS KAKENH (grant number JP17K15694). Work in the SCB unit at OIST is funded by core subsidy from Okinawa Institute of Science and Technology Graduate University.

## Author contributions

B.S. conceived the study and performed the experiments. A.B. and B.S. designed and engineered the micron pillar arrays. H.C. contributed reagents. B.S. and U.S. analyzed the data. B.S. wrote the manuscript with input from all authors.

